# Evolutionary changes in germ granule mRNA content are driven by multiple mechanisms in *Drosophila*

**DOI:** 10.1101/2023.02.21.529147

**Authors:** Dominique A. Doyle, Florencia N. Burian, Benjamin Aharoni, Annabelle J. Klinder, Melissa M. Menzel, Gerard Carlo C. Nifras, Ahad L. Shabazz-Henry, Bianca Ulrich Palma, Gisselle A. Hidalgo, Christopher J. Sottolano, Bianca M. Ortega, Matthew G. Niepielko

## Abstract

The co-packaging of mRNAs into biomolecular condensates called germ granules is a conserved strategy to post-transcriptionally regulate mRNAs that function in germline development and maintenance. In *D. melanogaster*, mRNAs accumulate in germ granules by forming homotypic clusters, aggregates that contain multiple transcripts from a specific gene. Nucleated by Oskar (Osk), homotypic clusters in *D. melanogaster* are generated through a stochastic seeding and self-recruitment process that requires the 3’ UTR of germ granule mRNAs. Interestingly, the 3’ UTR belonging to germ granule mRNAs, such as *nanos* (*nos*), have considerable sequence variations among *Drosophila* species. Thus, we hypothesized that evolutionary changes in the 3’ UTR influences germ granule development. To test our hypothesis, we investigated the homotypic clustering of *nos* and *polar granule component* (*pgc*) in four *Drosophila* species and concluded that homotypic clustering is a conserved developmental process used to enrich germ granule mRNAs. Additionally, we discovered that the number of transcripts found in *nos* and/or *pgc* clusters could vary significantly among species. By integrating biological data with computational modeling, we determined that multiple mechanisms underlie naturally occurring germ granule diversity, including changes in *nos*, *pgc*, *osk* levels, and/or homotypic clustering efficacy. Finally, we found that the *nos* 3’ UTR from different species can alter the efficacy of *nos* homotypic clustering, resulting in germ granules with reduced *nos* accumulation. Our findings highlight the impact that evolution has on the development of germ granules and may provide insight into processes that modify the content of other classes of biomolecular condensates.

## Introduction

Throughout the animal kingdom, germline function and maintenance require the formation of ribonucleoprotein (RNP) granules called germ granules [1–7]. Specifically, germ granules are biomolecular condensates that function in the post-transcriptional regulation of mRNAs that have critical roles in germline differentiation, proliferation, and function [1–10]. In *Drosophila*, germ granules develop within a highly specialized cytoplasm called the germ plasm that develops at the posterior oocyte and persists in the early embryo for ~1-2 hours post fertilization [11, 12]. The formation of the germ plasm and its germ granules begins when *oskar* mRNA (*osk*) localizes and accumulates in the posterior of the oocyte, where it is translated and recruits additional proteins such as Vasa (Vas) and Tudor (Tud) to form germ granule protein ensembles. Although several proteins are known to comprise the germ granule protein ensemble, Osk is the only protein that is both necessary and sufficient for germ plasm formation [13–16]. Comprising the mRNA portion of germ granules are transcripts such as *germ-cell less* (*gcl*), *cyclinB* (*cycB*), *nanos* (*nos*), and *polar granule component* (*pgc*) [17, 18]. These genes are maternally transcribed by specialized cells called nurse cells, which then deposit the mRNAs into the oocyte’s bulk cytoplasm as RNPs containing single transcripts (Fig. 1) [17]. Simultaneously with the Osk driven formation of the germ granule protein ensembles, single transcript RNPs that are diffusing through the bulk cytoplasm localize to the germ plasm by incorporating into germ granules through a stochastic seeding and self-recruitment process that results in the formation of homotypic clusters, mRNA aggregates that contain several copies of the same transcript (Fig. 1) [17, 19, 20]. The formation and growth of germ granules occurs continuously for ~19 hours from oocyte stages (~9 to 14) to the early embryo, when the degradation of *osk* mRNA has completed [15, 19–21].

**Figure 1:**
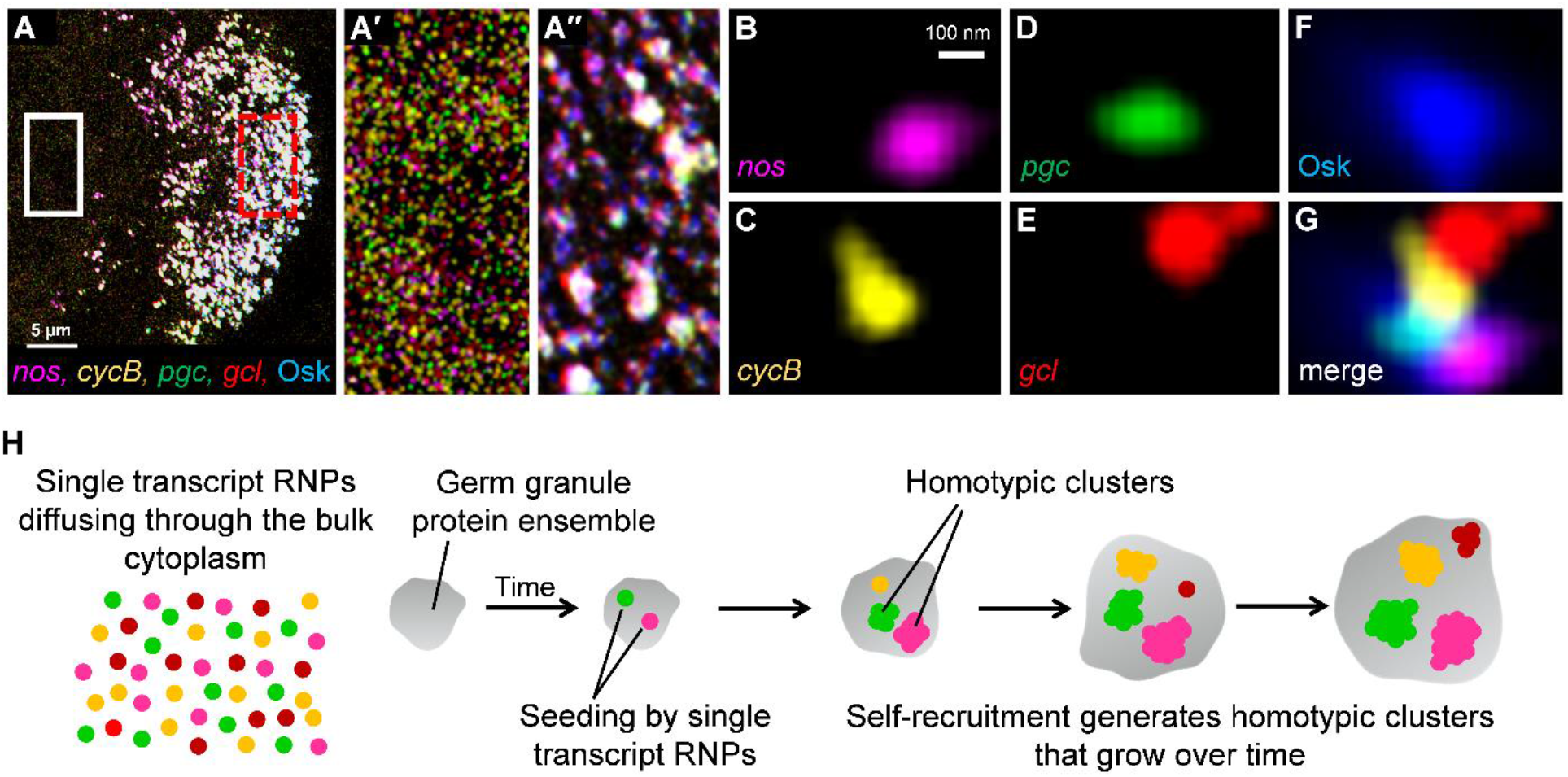
Localization of mRNAs to the germ plasm by forming homotypic clusters within germ granules in *D. melanogaster*. **A)** Super resolution max project image of germ plasm mRNAs *nos*, *cycB, pgc*, and *gcl* that are labeled using smFISH and Osk protein tagged with GFP in a stage 13 oocyte, posterior is the right. **A’)** Enlarged region of A marked by the white box that depicts the single transcript RNPs that are diffusing through the bulk cytoplasm of the oocyte. **A’’)** Enlarged region of germ granules (marked by Osk-GFP, blue), in the posterior germ plasm that is marked by the broken red box in A. **B-G)** Germ plasm mRNAs that have accumulated within a single germ granule by forming homotypic clusters. **H)** Overview of homotypic cluster formation in *D. melanogaster*.

Recent studies have employed a combination of techniques such as single molecule fluorescent *in situ* hybridization (smFISH), super-resolution microscopy, quantitative image analysis, and computational modeling to investigate germ granule development by quantifying the mRNA content of germ granules in *Drosophila melanogaster* (*D. mel*) [5, 6, 17, 19, 20, 22, 23]. These interdisciplinary methods have enabled the quantification of the number of transcripts found within a homotypic cluster (referred to as cluster size), the characterization of homotypic cluster size distributions for thousands of clusters found throughout a germ plasm (referred to as germ plasm landscape), the measuring of the frequency at which different types of homotypic clusters populate the same granule (referred to as co-localization), and the description of the relationship between the sizes of different homotypic clusters that are detected within the same granule [17, 19, 20]. Such features of germ granules can be collectively visualized through the Granule Census, an image analysis pipeline that transforms 3D confocal images of smFISH data into a quantified 2D quantified matrix, which can also be produced *in silico* using a computational model that simulates the formation of germ granules [19, 20].

The mRNA composition of germ granules relies primarily on three known mechanisms that work in conjunction with each other 1) the transcript levels of a particular germ granule mRNA, 2) the amount of *osk* present in the germ plasm, which controls the granule’s mRNA carrying capacity, and 3) the efficacy by which an mRNA can accumulate within a homotypic cluster, referred as the clustering factor, which has been shown to be influenced by *cis*-acting sequences called “clustering elements” found in the 3’ UTR [20, 22]. Although germ granule formation and mRNA composition has been described and modeled in *D. mel* and several mutants [17, 19, 20], it is unclear how conserved the germ granule formation process is in other *Drosophila* species, whether mRNA composition varies among species, and what germ granule assembly mechanisms can be influenced by evolutionary changes. Here, our goal is to investigate the evolution and development of *Drosophila* germ granules to gain insight into their diversity and to identify developmental mechanisms that are susceptible to modifications.

Previous studies have demonstrated that the accumulation of *nos* in the germ plasm of *D. mel* can be achieved using the *nos* 3’ UTR from other species despite considerable sequence variations identified in the *nos* 3’ UTR, suggesting conserved mechanisms that participate in germ granule formation [24]. However, the germ granule assembly process has not been explored in other *Drosophila* species, thus questions surrounding the impact that evolution has on germ granule assembly mechanisms and germ granule mRNA compositions have not been investigated. In this study, we examined the germ granule assembly process in four *Drosophila* species to identify both conserved and diverse features of *Drosophila* germ granules. We performed smFISH against *nos* and *pgc*, and carried out quantitative image analysis in *D. mel, D. virilis*, *D. pseudoobscura*, and *D. nebulsa* in developmentally early, mid, and late germ plasms. We find that single transcripts diffusing outside of the germ plasm is a conserved feature during the germ granule assembly process. Interestingly, we find that the germ granule mRNA composition can be highly diverse among species, including significant variation in cluster sizes of *nos* and/or *pgc* and co-localization rates between different homotypic clusters. By developing the Granule Content Transformation Analysis pipeline, we also determined that the overall mRNA composition of *D. nebulosa* was most divergent from all the other species. To identify which mechanism(s) generate diversity in germ granule mRNA compositions, we integrated qPCR data with computational modeling and found that changes to a combination of mechanisms, including transcript levels of *pgc*, *nos*, *osk* and/or clustering factor, contribute to germ granule diversity. To validate the importance of clustering factor in generating evolutionary changes in germ granule mRNA composition, we expressed the *nos* 3’ UTRs from *D. pseudoobscura* and *D. nebulasa* in *D. mel* and recapitulated expected reduction in the sizes of *nos* homotypic clusters. Together, our findings reveal the conserved and diverse features of *Drosophila* germ granules and demonstrate how germ granule mRNA content is subject to evolutionary changes. Additionally, we discovered multiple genetic mechanisms that are prone to evolutionary modifications that underlie *Drosophila* germ granules diversity. More broadly, our findings may offer insight into the evolution and development of other types of biomolecular condensates.

## Results

### Homotypic cluster formation is a conserved process in *Drosophila* species

To begin investigating the conservation of germ granule assembly in genus *Drosophila*, we explored 1) whether single RNP molecules of *nos* and *pgc* diffuse in the oocyte bulk cytoplasm and 2) whether each mRNA type generates homotypic clusters in the posterior germ plasm in other *Drosophila* species. We preformed *nos* and *pgc* smFISH experiments, confocal microscopy, and quantitative image analysis on stage 13 oocytes in *D. mel, D. pseudoobscura* (*D. pse*), *D. virilis* (*D. vir*), and *D. nebulosa* (*D. neb*). Stage 13 oocytes were chosen based on the presence of both single transcripts in the bulk cytoplasm and large homotypic clusters of *nos* and *pgc* in the posterior germ plasm in *D. mel* [19]. Similar to *D. mel*, we discovered that RNPs containing single *nos* or *pgc* transcript diffuse through the bulk cytoplasm, outside of the posterior germ plasm in all non-*D. mel* species tested (Fig. 2 and Supp Fig. 1A-B, see Material and Methods). To identify whether homotypic clustering exists in non-*D. mel* species, we performed quantitative image analysis to calculate how many transcripts reside in the brighter RNPs that were observed in the posterior germ plasm of each species (Fig. 2). In *D. mel*, we calculated the average number of transcripts in posterior *nos* RNP*s* was 7.50 ± 0.45 and for *pgc* RNPs it was 5.14 ± 0.27 which are both expected values based on previously published work (Supp Fig. 1C,D) [19]. For, *D. pse*, the average number of transcripts in posterior *nos* RNPs was 4.04 ± 0.34 and for *pgc* RNPs was 5.72 ± 0.98. For *D. vir*, the average number of transcripts in posterior *nos* RNPs was 3.23 ± 0.39 and for *pgc* RNP*s* was 2.74 ± 0.15. In *D. neb*, the average number of transcripts in the posterior *nos* RNP*s* was calculated to be 5.32 ± 0.34 and for *pgc* RNP*s* was 1.49 ± 0.05 (Supp Fig. 1C,D). In all species, larger *nos* and *pgc* RNPs were also detected (Supp Fig. 1E,F). Next, we analyzed the frequency that germ plasm *nos* RNPs and *pgc* RNPs reside within the germ granule by calculating their co-localization rates in each species. The co-localization for *D. mel* was 46.38% ± 2.32, which is similar to the expected previously published rate [19]. In *D. pse, D. vir*, and *D. neb*, the co-localization rate between germ plasm *nos* and *pgc* RNPs was 47.06% ± 3.77, 22.00% ± 3.54, and 18.38% ± 4.46 (Supp Fig. 1G), respectively, and were all greater than the previously published 10% rate for random co-localization [19]. In all species tested, the average number of transcripts that reside in germ plasm *nos* RNPs was greater than 3 while the co-localization rate between different mRNA types was under 50%. Furthermore, larger *nos* and *pgc* RNPs were observed throughout the germ plasm in all species tested (Supp Fig. 1). Together, these data demonstrate that transcripts from *nos* are more likely to co-localize with itself rather than co-localize with *pgc*, supporting a conserved process to localize mRNAs to the germ plasm through the formation of homotypic clusters within germ granules. Thus, we conclude that similar to *D. mel*, germ granules in other *Drosophila* species are heterogenous with respect to the types of homotypic clusters that they contain and the number of transcripts found in a homotypic cluster.

**Figure 2:**
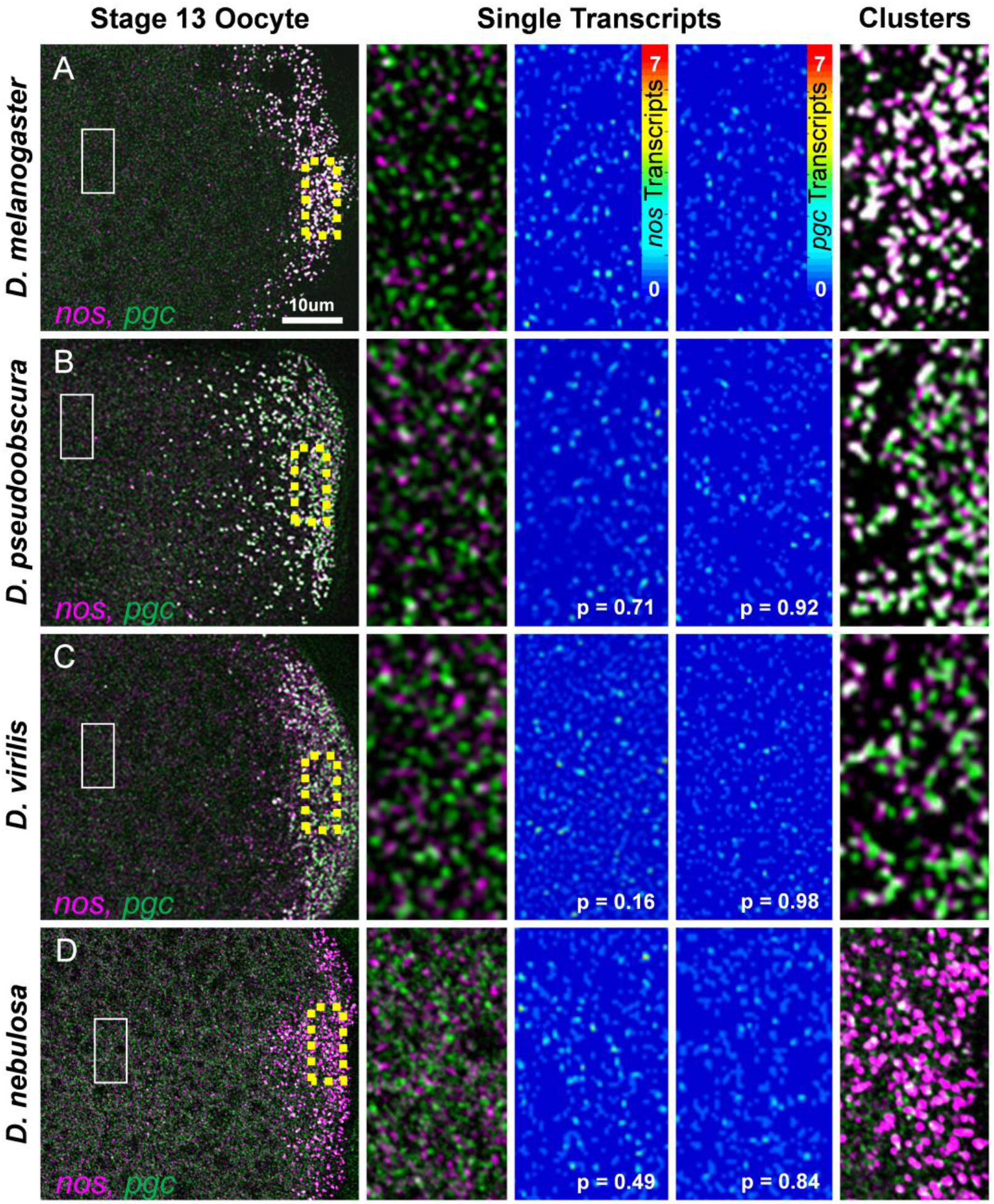
Single transcript RNPs in the bulk cytoplasm and posterior accumulation of *nos* and *pgc*. **Rows A-D, Column I**) Confocal images (max projection) of *nos* (magenta) and *pgc* (green) using smFISH in stage 13 oocytes. **A)** *D. melanogaster*, **B)** *D. pseudoobscura*, **C)** *D. virilis*, and **D)** *D. nebulosa* (posterior is to the right). **Column II)** Single transcripts that are not in germ granules and are dispersed throughout the bulk cytoplasm (white box in Column I). **Column III)** Heat maps of single *nos* molecules in the bulk cytoplasm. **Column IV)** Heat maps of single *pgc* molecules in the bulk cytoplasm, p-values between single molecule intensities from each species and *D. mel* show no difference. **Column V)** Posterior accumulation of *nos* and *pgc* in the germ plasm (yellow boxes in Column I). Images are a representation of at least 3 stage 13 germ plasms for each species.

### The mRNA composition of germ granules is diverse among different *Drosophila* species

Although we identified conserved germ granule features in other *Drosophila* species, such as the presence of single transcript RNPs in the bulk oocyte cytoplasm and homotypic clusters in the posterior germ plasm (Fig. 2 and Supp Fig. 1), we next probed whether germ granule mRNA compositions were diverse among *Drosophila* species by performing a Granule Census analysis on stage 13 oocytes in *D. mel, D. pse, D. vir*, and *D. neb*. The Granule Census transforms 3D confocal images of germ plasm *nos* and *pgc* smFISH data in a 2D quantified matrix to visualize the mRNA composition of germ granules within the germ plasm [19, 20]. The Granule Census revealed striking differences between the *nos* and *pgc* compositions of germ granules from *Drosophila* species. Specifically, we found that the average cluster size of *nos* was significantly smaller in all species tested when compared to *D. mel* p < 0.002. Additionally, *D. vir* and *D. neb* had significantly smaller clusters of *pgc* when compared to *D. mel* (p < 0.002) while *D. pse* produced similar sized *pgc* homotypic clusters as *D. mel* (p = 0.650) (Supp Fig. 1 and Fig. 3). In summary, we find that *vir* had the smallest *nos* cluster sizes while *D. neb* had the weakest germ granule accumulation of *pgc* when compared to *D. mel*. Furthermore, we determined that the co-localization rates between *nos* and *pgc* clusters were significantly lower in *D. vir* and *D. neb* (p < 0.001) when compared to *D. mel* (Supp Fig. 1G). Next, we analyzed the relationship between the sizes of co-localized *nos* and *pgc* clusters by calculating the slope of the best-fit-line of co-localized cluster sizes in different *Drosophila* species. For *D. mel*, the slope was 0.50 ± 0.02 and consistent with previously published results [19]. In *D. pse*, the slope increased to 1.24 ± 0.07 (p < 0.001) when compared to *D. mel*. For *D. vir* and *D. neb*, the slope was reduced to 0.33 ± 0.05 and 0.044 ± 0.0 (p = 0.032 and p < 0.001), respectively, when compared to *D. mel* (Fig. 3E). Together, these data demonstrate that homotypic clustering, co-localization rates, and the balance between the sizes of clusters that populate the same granule can be diverse or similar among different *Drosophila* species.

**Figure 3:**
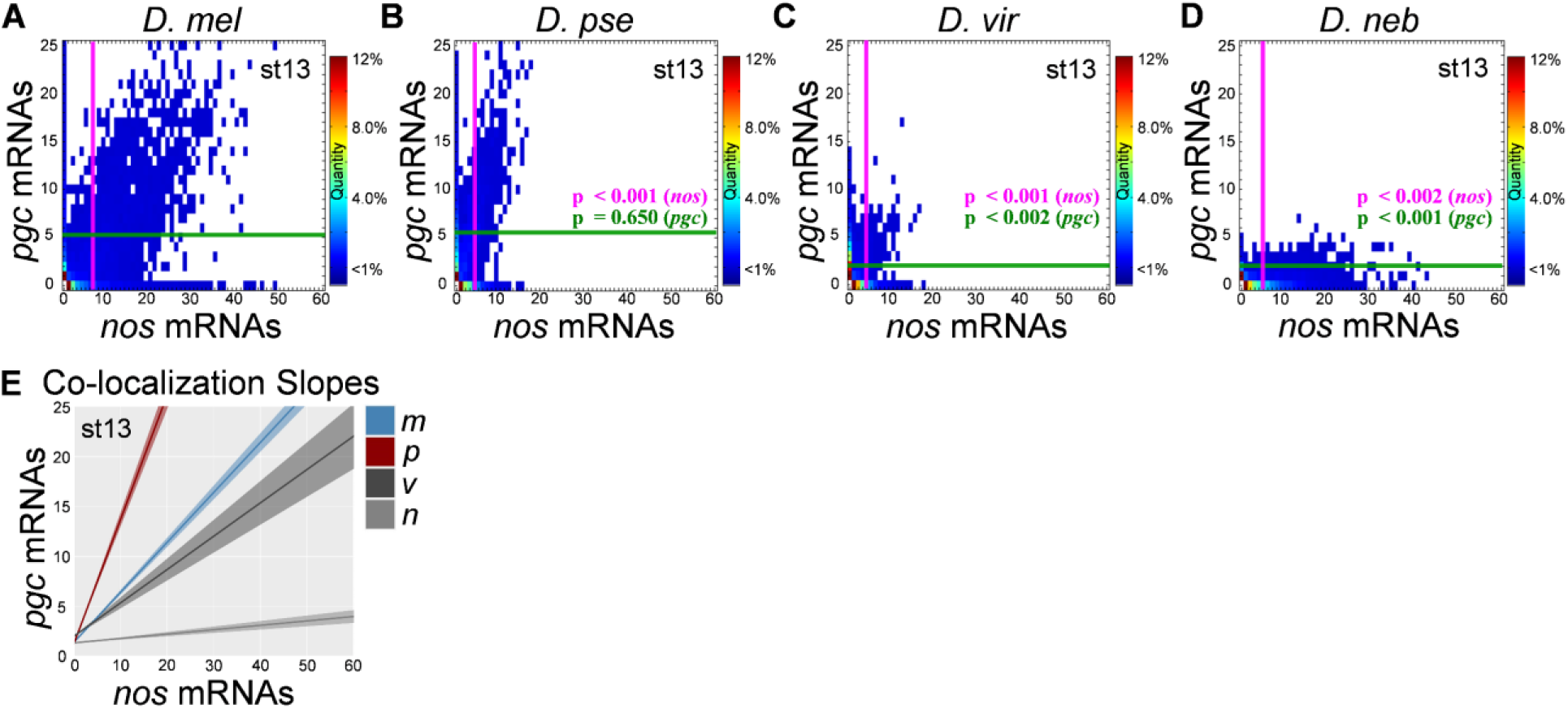
Germ granule censuses produced from various *Drosophila* species. **A-D)** The Granule Censuses produced from analyzing *nos* and *pgc* homotypic clusters in stage 13 germ plasms from *D. melanogaster* (*D. mel*), *D. psuedoobscura* (*D. pse*), *D. virilis* (*D. vir*), and *D. nebulosa* (*D. neb*). The magenta vertical line represents the average *nos* cluster size observed in the germ plasm while the green horizontal line represents the average *pgc* cluster size in the germ plasm in each census. **E)** Lines of best fit generated by plotting the cluster sizes of *nos* and *pgc* that reside within the granule (co-localized) for each species. The shaded regions represent the ± S.E.M for at least 3 biological replicates. The heatmaps in A-D represent the percent of granules that have a particular *nos* and *pgc* mRNA composition. For all species n > 5,000 homotypic clusters were analyzed and p-values were calculated based on comparing each species’ average *nos* or *pgc* clusters size to *D. mel’s* average *nos* or *pgc* cluster size.

Next, we aimed to characterize and score the overall change in the *nos* and *pgc* composition of germ granules between *Drosophila* species. To identify overall change, we developed a pipeline called the Granule Content Transformation Analysis (GCTAnalysis), which highlights the transformation from one germ plasm to another and scores the magnitude of change between two censuses where a larger value refers to a greater transformation (Fig. 4A). By calculating GCTAnalysis scores for each species pair, we found thar *D. mel* to *D. pse* had a score of 4.14, the score for *D. mel* to *D. vir* was 10.47, *D. mel* to *D. neb* was 13.31, *D. pse* to *D. neb* was 13.51, *D. vir* to *D. pse* was 8.12, and *D. vir* to *D. neb* was 12.55 for stage 13 oocytes (Fig. 4B-G). Next, we generated an undirected graph to visualize the magnitude of change between species’ germ plasm and in summary, our analysis between *nos* and *pgc* mRNA compositions revealed that the germ plasm between *D. mel* to *D. pse* were most similar while *D. neb* was the most different when compared to the other species (Fig. 4H). Next, we analyzed the number of *nos* and *pgc* mRNA compositions that are similar or unique among species’ germ plasms. Overall, we found that *D. mel* had the most unique *nos* and *pgc* mRNA compositions found within germ granules at 305, followed by *D. pse* at 52 and *D. neb* at 33. We did not observe any *nos* and *pgc* mRNA compositions that were unique to *D. vir* in stage 13 oocytes (Fig. 4I). Altogether, our data and comprehensive analysis show that the mRNA composition of germ granules is subject to evolutionary changes that generate germ plasm diversity among *Drosophila* species.

**Figure 4:**
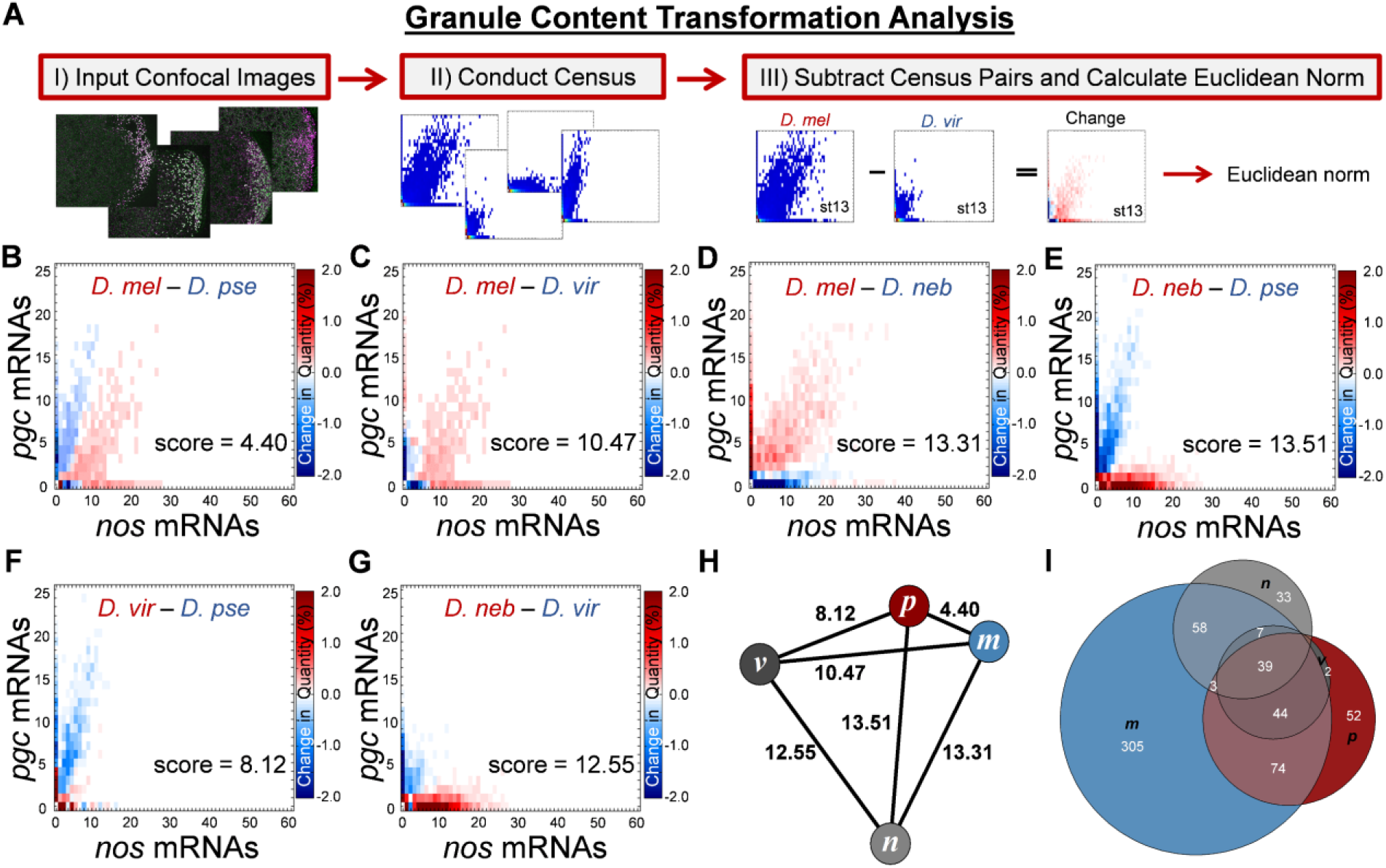
Scoring the overall diversity of *nos* and *pgc* germ granule content. **A)** Changes in germ granule mRNA content are scored using a pipeline called the Granule Content Transformation Analysis. The pipeline begins by acquiring 3D confocal images and transforming them into quantified 2D Granule Censuses. Next, a new census that represents a change between two Granule Censuses is generated by subtracting two censuses. To score the overall magnitude of change, the Euclidean norm of the resulting census is calculated. **B-G)** Changes in *nos* and *pgc* gem granule mRNA content between different pairs of *Drosophila* species in stage 13 oocytes with the overall score noted. The heat map represents change in quantity for a given *nos* and *pgc* combination found in the Granule Census, the darker the color (blue or red), the larger the change between the paired species, while white represents no change in a particular composition between the different species. The names of species are color coordinated based on mRNA composition that is found more often for that species. **H)** Undirected graph where nodes are represented by species and edges are represented by the score calculated in B-G, *melanogaster* (*m*), *pseudoobscura* (*p*), *virilis* (*v*), and *nebulosa* (*n*). **I)** Venn diagram that displays the number of *nos* and *pgc* mRNA compositions that are unique or overlapping between different *Drosophila* species.

### Presence of germ granule diversity in early stages of germ plasm formation

The germ plasm begins to form during oocyte stage 9 and continues through oocyte stage 14 and the early embryo, for a total development time of ~19 hours [20, 25]. Given the observed germ granule diversity observed among different *Drosophila* species during stage 13 oocytes (Figs. 3 and 4), we next examined whether diversity in germ granule mRNA composition can be observed early during germ plasm formation. To explore when germ granule diversity can be detected, we performed smFISH and the Granule Census for *nos* and *pgc* on stage 10 oocytes for *D. mel, D. pse, D. vir*, and *D. neb*. The average *nos* cluster sizes were 3.14 ± 0.11 for *D. mel*, 2.17 ± 0.09 for *D. pse*, 2.10 ± 0.09 for *D. vir*, and 3.34 ± 0.13 for *D. neb*. On average, *nos* cluster sizes were smaller by only ~1 transcript in *D. pse* and *D. vir* (p > 0.001) while *nos* cluster sizes were similar in *D. neb* when compared to *D. mel* (p = 0.35). For *pgc*, the average cluster size was 2.35 ± 0.17 in *D*. mel, which was not significantly different when compared to *D. pse* (2.59 ± 0.08) and in *D. vir* (2.17 ± 0.30), p > 0.71 for both. However, diversity in *pgc* homotypic clustering was observed in *D. neb* in that homotypic clusters for *pgc* were not readily detected, with an average cluster size of 1.59 ± 0.05, which was significantly different when compared to *D. mel*, p = 0.02 (Fig. 5). Next, we analyzed the co-localization rates between *nos* and *pgc* in stage 10 oocytes and found that *D. mel* had a co-localization rate of 29.12% ± 3.34, which was similar to previously published data [19]. This co-localization rate was not significantly different when compared to *D. pse* (29.00% ± 3.00), *D. vir* (25.00% ± 8.40), and *D. neb* (22.5% ± 2.90), p > 0.55 and were all greater than the 10% random co-localization rate [19]. Following the calculation of co-localization rates, we analyzed the slope generated by the sizes of co-localized *nos* and *pgc* homotypic clusters for all species in stage 10 oocytes and found that *D. mel* had a slope of 0.25 ± 0.03, which was similar to the slope generated by *D. vir* (0.32 ± 0.07 p = 0.63). The slope generated by co-localized stage 10 *nos* and *pgc* clusters in *D. pse* (0.73 ± 0.08) was significantly greater than *D. mel* (p < 0.001). The slope in *D. neb* stage 10 oocytes was 0.08 ± 0.02, which was significantly less than *D. mel*(p < 0.03), further demonstrating the presence of diversity in the balance between the number of *nos* and *pgc* transcripts between *D. mel* and other species (Fig. 5). Together, these data show that germ granule diversity in early germ plasm formation is detected for some germ granule features.

**Figure 5:**
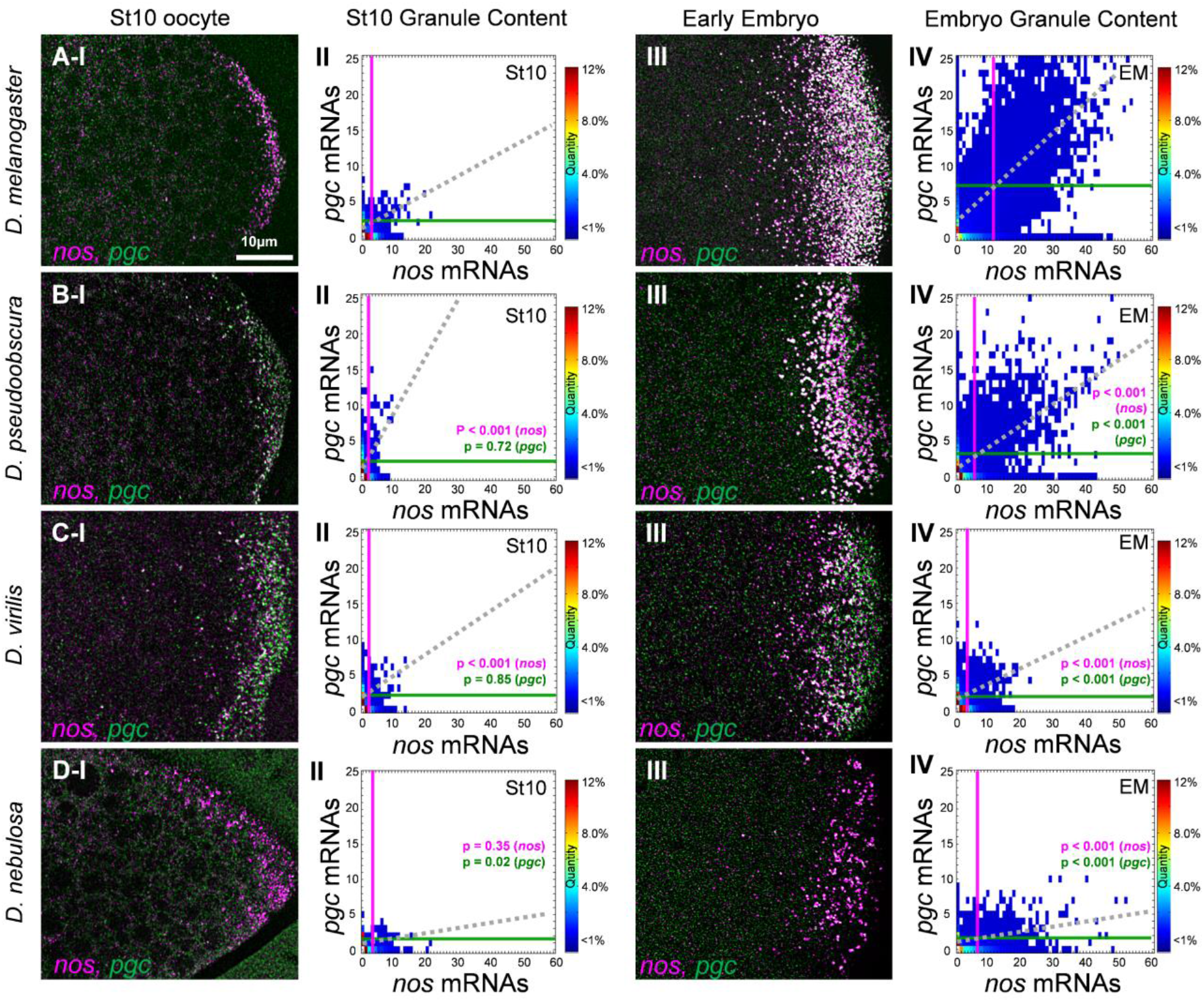
Germ granule mRNA composition at stage 10 and the early embryo in *Drosophila* species. **Row A)** Rows A-D, Column I) Confocal images (max projection) of *nos* (magenta) and *pgc* (green) using smFISH in stage 10 oocytes. **A)** *D. melanogaster*, **B)** *D. pseudoobscura*, **C)** *D. virilis*, and **D)** *D. nebulosa* (posterior is to the right). **Rows A-D, Column II)**, Granule Census for each species’ germ *plasm* in stage 10 oocytes. **Rows A-D, Column III)** Confocal images (max projection) of *nos* (magenta) and *pgc* (green) using smFISH in the early embryo. **Rows A-D, Column IV)** Granule Census for each species’ germ plasm in the early embryo. For all species, images are a representation of at least 3 stage 10 and early embryo germ plasms. Each stage 10 census includes n >1,300 granules while each early embryo census includes n >10,000 granules. The magenta vertical line represents the average *nos* cluster size observed in the germ plasm while the green horizontal line represents the average *pgc* cluster size in the germ plasm in each census. Dotted gray line represents the line of best fit that is produced by the sizes of co-localized *nos* and *pgc* clusters.

### Germ granule formation dynamics vary among *Drosophila* species

In *D. mel, nos* and *pgc* clusters persist in the germ granules of 1-2 hour old embryos [17]. Thus, we examined whether germ granule diversity is also observed in the early embryo. We first determined that the average germ granule content of *nos* in *D. mel* was 11.13 ± 0.75 and was similar to previously published data [19]. Next, we found that the average number of *nos* transcripts observed in germ granules for *D. pse* was 5.37 ± 0.14, for *D. vir* was 3.07 ± 0.27, and in *D. neb* was 6.50 ± 0.40, which were all significantly less than the average in *D. mel* (p < 0.001, Fig. 5). For *pgc*, the average germ granule content in *D. mel* germ granules in the early embryo was 6.56 ± 0.59 and was also similar to previously published data [19]. The average number of *pgc* transcripts observed in germ granules for *D. pse* was 3.41 ± 0.27, for *D. vir* was 2.23 ± 0.07, and in *D. neb* was 1.60 ± 0.19, which were all significantly less than the average when compared to *D. mel* (p < 0.001, Fig. 5). As for co-localization, we found that in *D. pse, nos* and *pgc* had a co-localization of 36.00% ± 3.53, in *D. vir* it was 28.70% ± 2.00, and in *D. neb* it was 22.71% ± 5.80. All these values were all significantly lower than the 56.50% ± 1.91 co-localization rate calculated for *D. mel* (p < 0.005). Analyzing the slopes created by the sizes of co-localized *nos* and *pgc* clusters, we found that *D. neb* had the smallest slope at 0.04 ± .02, followed by *D. vir* at 0.17 ± 0.03, and *D. pse* at 0.36 ± 0.08 (Fig. 5). In *D. vir* and *D. neb*, these values were all significantly lower than the 0.49 ± 0.04 slope measured for *D. mel* (p < 0.001), demonstrating that the balance between the number of *nos* and *pgc* transcripts can vary between *D. mel* and other species in the early embryo. Together, these data show that in addition to the oocyte germ plasm, diversity in *nos* and *pgc* clustering is also observed in the germ granules of early embryos.

In the early embryo, germ granules in *D. mel* contain more *nos* and *pgc* transcripts than what is observed in stage 13 oocyte germ plasm. Specifically, the average number of *nos* transcripts in *D. mel* germ granules increases from ~7 to ~11 from stage 13 oocytes to the early embryo while the average number of *pgc* transcripts increases from ~6 to ~7 transcripts, which supports a model of continuous mRNA accumulation for both *nos* and *pgc* [17, 19]. Our next goal was to explore whether other species have similar accumulation dynamics and to capture any changes in the behavior of germ granule formation. First, we compared the average cluster sizes of *nos* between stage 13 oocytes and the early embryo in non-*D. mel* species. In *D. pse* and *D. neb*, the observed increase in *nos* cluster size between stages was significant (p < 0.044), suggesting that like *D. mel*, these species also continue to accumulate *nos* in the early embryo, however the increase in *nos* for these species was modest in comparison to the change observed in *D. mel* (Figs. 3 and 5). Unlike the other three species, *nos* cluster sizes were static between stage 13 and the early embryo for *D. vir* at ~3 transcripts. As for *pgc*, cluster sizes were static between stage 13 and the early embryo for *D. neb* at ~1.5 transcripts and *D. vir* at ~2 transcripts, suggesting that unlike *D. mel, pgc* and/or *nos* accumulation does not continue in the early embryo for these species. Interestingly, a small decrease in *pgc* cluster size was detected from stage 13 and early embryo in *D. pse* (p > 0.025), suggesting possible degradation of *pgc* homotypic clusters in the early embryo in *D. pse* (See Discussion).

To measure and visualize overall differences in *nos* and *pgc* accumulation dynamics between each species, we calculated and triangulated GCTAnalysis scores between stage 10, 13, and the early embryo for each species and analyzed the changes in triangle shapes (Supp Fig. 2). In *D. mel*, the scores generated an obtuse scalene triangle, with the obtuse angle of 146° at the stage 13 node. However, in *D. pse* and *D. vir*, the scores generated acute scalene triangles with angles of 81° and 88° degree at the stage 13 node, respectively. In *D. neb*, the scores also created an obtuse scalene triangle, however the obtuse angle of 160° was at the stage EM node, unlike the obtuse angle at the stage 13 node in *D. mel* (Supp Fig. 2). The triangles highlight differences in germ granule formation dynamics such as the lack of change from stage 13 to the embryo in *D. vir* (Supp Fig. 2C) and a regression of germ granule composition from stage 13 towards stage 10 in *D. neb* embryos (Supp Fig. 2D). By generating species specific triangles, we were able to visualize how the developmental dynamics of germ granules vary between stages and among species. Together, these data show diversity in the accumulation of *nos* and *pgc* transcripts in the early embryo along with variability in the overall developmental dynamics of germ granules.

### Computational modeling recapitulates germ granule diversity

Our results highlight how germ granules mRNA compositions of *nos* and *pgc* homotypic clusters vary among different *Drosophila* species (Fig. 2–5), thus we next aimed to understand the mechanisms that contribute to such evolutionary changes. In *D. mel*, the germ granule accumulation of *nos* and *pgc* can be altered by changing the transcript expression levels of *nos, pgc*, and/or *osk*. Furthermore, *cis-*acting elements found in the 3’ UTR also contributes to a gene’s specific “clustering factor,” which represent the efficacy by which a gene’s transcript can form a homotypic cluster within a germ granule [19, 20, 22]. Thus, we hypothesized that the diversity observed in germ granule mRNA composition among *Drosophila* species (Figs. 3 – 5) is driven by one or a combination of these mechanisms. To test our hypothesis, we first measured the transcript levels of *nos*, *pgc*, and *osk* in *D. pse, D. vir*, and *D. neb* and compared each expression to their *D. mel* counterpart (See Material and Methods). Relative to *D. mel*, we found that *osk* levels were not significantly different in *D. neb* (p > 0.13). However, *D. pse osk* levels were higher at 2.00 ± 0.23 (p < 0.0001), while *osk* levels in *D. vir* was lower at 0.43 ± 0.10 (p < 0.003) (Supp Fig. 3). For *nos* expression levels, *D. pse* and *D. neb* did not show a significant change in expression levels p > 0.50 and only *D. vir* had a significantly lower expression level (0.65 ± 0.09, p < 0.02) when compared to *D. mel*. Like *nos* expression, *pgc* in *D. pse* and *D. neb* did not show a significantly change in expression levels p = 0.99 and only *D. vir* had a lower expression level (0.47 ± 0.08, p < 0.008) when compared to *D. mel* (Supp Fig. 3). In *D. vir*, we reason that the reduction in both *nos* and *pgc* cluster sizes, when compared to *D. mel*, corresponds to the observed decrease in both *nos*, *pgc*, and *osk* levels (Figs. 3 and 5 and Supp Fig. 3). However, in *D. pse, nos* cluster sizes are reduced despite an increase in *osk* levels and equal level expression of *nos*. Furthermore, in *D. neb* where the average *nos* clusters are smaller than *D. mel’s* and poor clustering of *pgc* is observed, there are no changes in *nos*, *pgc*, or *osk* levels (Figs. 3 and 5 and Supp Fig. 3). These data suggest that changes in expression levels alone cannot account for the evolutionary changes in germ granule mRNA composition for every species and therefore, we hypothesized that differences in the clustering factor also contribute to germ granule diversity in *D. pse* and *D. neb*.

To measure how changes in clustering factor contribute to the observed germ diversity found in *D. pse* and *D. neb*, we combined computational modeling with our expression level data and calculated the clustering factor of *nos* and *pgc* in non-*D. mel* species [20]. Specifically, we adjusted the model’s transcript pool parameters to the relative expression levels of *nos* and *pgc* and set the germ granule’s carrying capacity (cc) parameter to the relative *osk* level from each species. To predict clustering factors for *nos* and *pgc* in *D. pse* and *D. neb*, we fitted the average cluster size to a clustering factor standard curve (Supp Fig. 4A,B and see Material and Methods). We found that in *D. pse*, the *nos* and *pgc* clustering factors are 0.20 and 0.14, respectively. In *D. neb*, we calculated the *nos* and *pgc* clustering factors to be 0.46 and 0.14, respectively (Supp Fig. 4A,B). In both cases, the clustering factors are smaller than the previously determined values of 0.74 for *nos* and 0.48 for *pgc* in *D. mel* [20]. Next, we generated modeling profiles for each species where the parameters are represented by the observed expression levels and the calculated clustering factors (Fig. 6A). Using the modeling profiles, we adjusted the model to recreate each species’ biological conditions *in silico* and simulated germ granule formation that recapitulated the Granule Census in the early embryo (Fig. 6). Using the model’s accuracy score [20], we determined that *D. vir’s* model was 91% accurate, *D. pse* model’s was 88% accurate, and *D. neb’ s* model was 80% accurate, when compared to their biological counterparts. Together, our biological and modeling data provide system level evidence that multiple and combined mechanisms contribute to the evolution and development of *nos* and *pgc* compositions in *Drosophila* germ granules.

**Figure 6:**
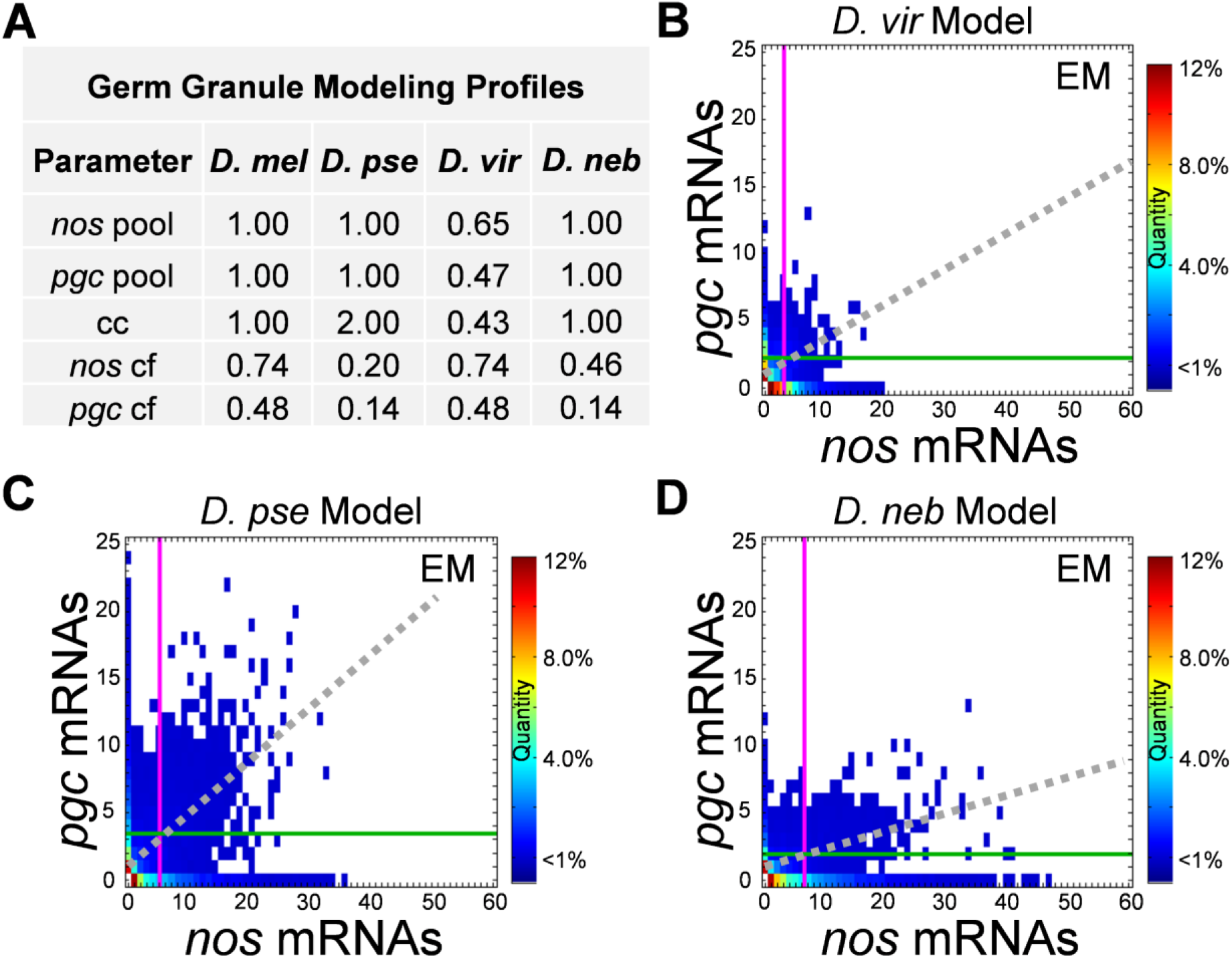
Computational modeling recapitulates germ granule diversity in the early embryo. **A)** Germ granule modeling profiles for each species used to generate embryonic germ granules *in silico*. **B-D)** Germ Granule Censuses produced using computational modeling from **B)** *D. vir*, **C)** *D. pse*, and **D)** *D. neb* modeling profile in panel A. The magenta vertical line represents the average *nos* cluster size observed in the germ plasm while the green horizontal line represents the average *pgc* cluster size in the germ plasm in each census. The dotted gray line represents the line of best fit produced by *nos* and *pgc* cluster sizes that reside within the same granule.

### Sequence diversity in the *nos* 3’ UTR yields differences in homotypic clustering

Although a combination of changes to the expression levels of *nos*, *pgc*, and *osk* can account for the observed mRNA compositions of germ granules in *D. vir* (Fig. 6A, B), our computational modeling predicts that a reduction in clustering factor, which represents homotypic clustering efficacy, contributes to the observed mRNA compositions of germ granules in *D. pse* and *D. neb* (Fig. 6). In *D. mel*, the clustering factor for *nos* is regulated, in-part, by sequences found in the 3’ UTR [20, 22]. Interestingly, aligning the *nos* 3’ UTRs between *D. mel, D. pse*, and *D. neb* revealed considerable sequence variation resulting in evolutionary distances of 1.06 between *D. neb* and *D. mel*, 0.67 between *D. pse* and *D. mel*, and 0.88 between *D. neb* and *D. pse* (see Materials and Methods). Thus, we hypothesized that differences within the *nos* 3’ UTR can alter *nos* homotypic clustering by reducing *nos’s* clustering factor (Fig. 6A). To test our hypothesis, we generated *D. mel* flies to have *nos* expressed with the 3’ UTR of either *D. pse* (*D. mel ^pse nos^* ^3’ UTR^) or *D. neb* (*D. mel ^neb nos^* ^3’ UTR^) and investigated whether each 3’ UTR could 1) rescue *nos* localization to the germ plasm by forming homotypic clusters and 2) reduce *nos* cluster sizes by lowering *nos’s* clustering factor. To answer these questions, we preformed smFISH, confocal microscopy, and quantitative image analysis on *nos* in the early embryos collected from *D. mel ^pse nos^* ^3’ UTR^ and *D. mel ^neb nos^* ^3’ UTR^ flies. We found that the *nos* 3’ UTR from *D. pse* and *D. neb* rescues the localization of *nos* to the posterior germ plasm by forming homotypic clusters (Fig. 7). Next, we calculated the average cluster sizes of *nos* in both genotypes and compared them to wild-type *nos*. In *D. mel ^pse nos^* ^3’ UTR^ and *D. mel ^neb nos^* ^3’ UTR^ the average cluster sizes were 2.66 ± 0.23 and 3.45 ± 0.83, respectively. These cluster sizes were significantly different from wild-type *D. mel* which had an average *nos* cluster size of 11.16 ± 0.76, p < 0.0001. Next, we measured the relative expression levels of chimeric *nos* transcripts and found that the *nos* in *D. mel ^neb nos^* ^3’ UTR^ had equal expression to that of wild type *D. mel*, p = 0.94. In *D. mel ^pse nos^* ^3’ UTR^, we found that the *nos* transcripts levels were reduced to 0.724 when compared to wild-type *D. mel*, p = 0.039 (Supp Fig. 4C). Although we observe 27.6% less *nos* transcripts in *D. mel ^pse nos^* ^3’ UTR^ flies, this decrease cannot account for the ~4-fold decrease in average cluster size since 37% less *nos* expression results in only ~35% fewer *nos* transcripts in the granule [19, 20]. Thus, we hypothesized that the *nos* 3’ UTRs from different *Drosophila* species reduced cluster sizes by reducing *nos’s* clustering factor. To test our hypothesis, we calculated the clustering factor of the *D. neb nos* 3’ UTR and *D. pse nos* 3’ UTR in *D. mel* by fitting the average cluster sizes to a standard curve generated by the germ granule computational model (Supp Fig. 4D,E). We determined that the clustering factor for the *D. neb nos* 3’ UTR was 0.29 while the clustering factor for *D. pse nos* 3’ UTR was 0.24 when expressed in *D. mel*. In both cases, the clustering factor is reduced when compared the wild-type 0.74 value [20]. Next, we measured the impact that clustering factor has on the overall localization of *nos* to the germ plasm by comparing the distribution of *nos* cluster sizes between wild-type *D. mel* and *D. mel ^neb nos^* ^3’ UTR^ and finding that the two distributions only have 38% overlap (Supp Fig. 4F). Our combined modeling and experimental data demonstrate that sequence variation within the *nos* 3’ UTR can diversify *nos* homotypic clustering, indicating that the evolution of noncoding sequences influences germ granule development.

**Figure 7:**
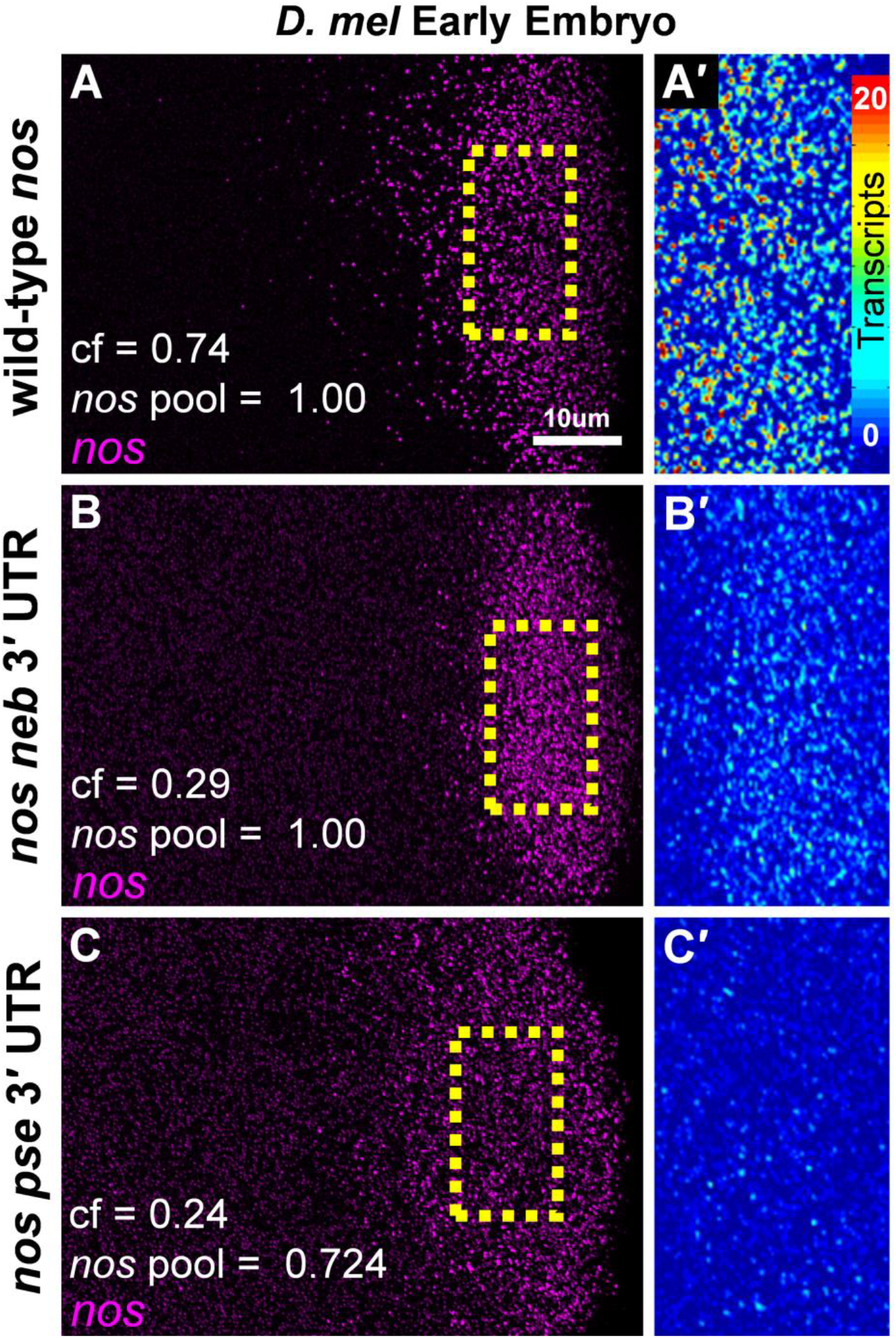
Evolutionary changes in the *nos 3* UTR influence the efficacy of *nos* homotypic clustering. **A-C)** Confocal images (max projection) of *nos* (magenta) using smFISH in the early embryo germ plasm. **A)** Wild-type *D. mel*, **B)** *D. mel* with the *nos* 3’ UTR from *D. neb* (*nos neb* 3’ UTR), and **C)** *D. mel* with the *nos* 3’ UTR from *D. pse* (*nos pse* 3’ UTR). Images are a representation of at least 4 germ plasms. Clustering factor values (cf) and *nos* expression levels (*nos* pool) for each genotype are reported in each panel. **A’ - C’)** Enlarged regions of the broken yellow boxes in A-C shown as heatmaps set to the same scale.

## Discussion

Although it is known that *nos* localizes to the posterior germ plasm in other *Drosophila* species such as *D. vir* [26], the analysis of *Drosophila* germ granule composition and assembly have been limited to a single species, *D. mel* [17–20, 22, 23]. Despite those studies providing much needed insight into the components, composition, and mechanisms that underlie germ granule formation, questions such as to what extent are those features conserved and how has the developmental process changed throughout evolution can only be answered by surveying additional species. Recent analysis of the germ plasm from zebrafish showed that similar to *D. mel*, homotypic clusters of *nanos3* and other mRNAs are present and their formation also requires a master organizing protein [26, 27]. The appearance of homotypic clustering across phyla introduces intriguing questions regarding the impact that evolution has on germ granule content and the mechanisms that influence their assembly and compositions. Using genus *Drosophila*, we address these evolutionary questions by providing a comprehensive cross-species analysis that explores the conserved and diverse features of germ granules. In doing so, we revealed how evolution impacts germ granule features while identifying genetic mechanisms in the assembly process that are susceptible to modifications.

The mRNA from an estimated 61 genes localize to the germ plasm in *D. mel* [11], including four that have been evaluated for homotypic clustering, *nos*, *pgc, gcl* and *cycB* (Fig. 1) [6]. Here, we focused on only two, *nos* and *pgc*, since clustering factor have been previously described for both mRNAs types [20, 22]. Given the extensive diversification of germ granules with respect to *nos* and/or *pgc* content that we report in this work, a broader sense of how germ granule mRNA content evolves should be obtained by studying additional germ granule mRNAs in future cross-species studies. In *D. mel*, the mRNA of the master organizer of germ granules, *osk*, also forms homotypic clusters [17]. However, this occurs in different germ plasm condensates called founder granules which are mutually exclusive with germ granules [21]. Here, our initial interspecies germ plasm study only focuses on the homotypic clustering of germ granule mRNAs. Whether diversification occurs in founder granules that connects to changes in germ granules has not been tested and should be explored in future germ plasm studies. In *D. mel*, a sub-granule organization for homotypic clusters has been described in that larger clusters tend to be located centrally within the granule. Specifically, *nos* clusters tend to be located more centrally than *pgc* clusters in *D. mel*, given the average size differential between the two cluster types [6, 18]. Although no function has yet been identified for this sub-granule organization [18], our results where a species’ *nos* and *pgc* clusters tend to be of similar size (Supp Fig. 1C,B and Fig. 5) raises interesting questions regarding the conservation of germ granule sub-cellular organization and whether such organization can exist in species where there is little to no cluster size differential among homotypic clusters. Further interspecies analysis into the conservation of sub-cellular organization of homotypic clusters may provide additional insight into the functional importance of sub-granule architecture. Regardless of our study’s limitation to only two germ granule mRNAs, our findings open the door for exciting future research that focuses on the evolution and development of additional homotypic clusters, founder granules, and sub-granule configuration and function.

We found that in *D. vir*, the differences in germ granule composition, when compared to *D. mel*, is driven by a combination of reduced *nos*, *pgc*, and *osk* levels. In the computational model, we represent such differences by adjusting parameters that mimic the transcript pools of all three transcripts by reducing their transcription rates (Fig. 6). Thus, we reason that differences in homotypic clustering can be attributed to variations at the transcriptional level. Alternatively, primordial germ cell accumulation of mRNAs in *D. mel* relies on mechanisms that protect mRNA from maternal degradation [28]. Thus, this study cannot rule out that differences in maternal degradation alone or in combination with reduced transcription rates contribute to the disparities in transcript levels among species. In *D. pse* and *D. neb*, we found that the *nos* 3’ UTR is reducing *nos’s* ability to accumulate within germ granules, as measured by the clustering factor value (Figs. 6 and 7). Although our modeling results successfully predicted a decrease in clustering value in *D. pse* and *D. neb*, the *nos* 3’ UTR in *D. neb* was predicted to be 0.46, which was higher than the 0.29 value determined when its 3’ UTR was expressed in *D. mel* (Figs. 6 and 7). In *D. mel*, the germ granule protein ensemble contains proteins such Osk, Vas, and Tud [13–16]. Furthermore, *nos* has been shown to bind to the Lotus domain of Osk in *D. mel* [29]. Thus, it is possible that germ granule proteins may also contribute to the clustering factor value and that co-evolution between 3’ UTRs and germ granule proteins has occurred, thus generating discrepancies between predicted and experimental clustering values that were calculated when 3’ UTRs were expressed in different species. Providing support for the co-evolution between 3’ UTRs and germ granule proteins is the finding that Osk from *D. vir* cannot concentrate enough germ cell mRNA determinants at the posterior to rescue primordial germ cell formation in *D. mel* [30]. In other systems, only a single mutation within the RNA-binding protein fused in sarcoma (FUS) was sufficient to induce structural changes to its condensates [31], further providing evidence for a potential role for changes in RNA-binding proteins contributing to altered condensate structures. In this study, we focus on the mRNA portion of the germ granule and whether evolutionary changes within germ granule proteins are also contributing to the differences highlighted in this study cannot be ruled out. Understanding how the protein portion of germ granules are changing and whether there is co-evolution between specific 3’ UTR clustering elements and germ granule proteins should be addressed in follow up studies to gain additional insight into the evolution and development of biomolecular condensates.

In addition to the diversity in mRNA compositions that we discovered, we found that the overall dynamics of homotypic cluster growth vary among *Drosophila* species (Supp Fig. 2). In *D. vir*, we observed no significant growth in *nos* and/or *pgc* homotypic clusters, generating a mostly static germ plasm landscape from stage 13 to early embryo despite the detection of single transcripts outside of the germ plasm (Fig. 5 and Supp Fig. 2C). These data suggest that *D. vir* germ granules may reach their carrying capacity earlier than the other species and supports a model where *D. vir* germ granules have a smaller carrying capacity when compared to other species (Fig. 6). In *D. neb, pgc* homotypic clusters were not readily observed in stage 10 oocytes and only developed smaller clusters that do not grow from stage 13 to the early embryo (Fig. 5 and Supp Fig. 2C). Additionally, *pgc* clusters decrease in average cluster size in *D. pse* from stage 13 to early embryo (Figs. 3B and 5B). One possible scenario that could explain a decrease in *pgc* cluster sizes from stage 13 to the early embryo in *D. pse* is that it may be translated and degraded earlier during the maternal-to-zygotic transition. In *D. mel, pgc* functions in establishing transcriptional quiescence in primordial germ cells to help maintain their a germline fate [8, 32–34]. However, complete loss of *pgc* does not fully prevent the specification of primordial germ cells in *D. mel*, suggesting that *pgc* has an important, but not essential role in maintaining cell’s germline fate due to potentially overlapping or redundant mechanisms [32, 35]. Thus, small *pgc* cluster sizes that were observed in non-*D. mel* species could signify the presence of additional mechanisms to maintain transcriptional quiescence in primordial germ cells, and therefore, eliminating a reliance on the formation of larger *pgc* clusters in those species. Interestingly, supplying additional Nos protein to primordial germ cells has been shown to rescue primordial germ cell viability in *pgc* mutants [35]. Thus, differences in translational regulation could also serve as a mechanism to compensate for reduced abundance of *nos* and/or *pgc* within non-*D. mel* species to ensure primordial germ cell viability. Nonetheless, our study highlights differences in mRNA localization in the germ plasms across multiple *Drosophila* species, and whether there are redundancies and/or translational mechanisms that compensate for different germ granule compositions to ensure germline viability should be investigated in future germ plasm studies.

In genus *Drosophila, D. mel, D. pse*, and *D. neb* are categorized in subgenera *Sophophora*, with *D. mel* located in the *melanogaster* group, *D. neb* in the *willistoni* group, and *D. pse* in the *obscura* group. *D. vir* is organized within the *virilis* group located in subgenera *Drosophila* [36, 37]. Interestingly, our results show that the germ plasm landscape in the late oocyte is most divergent between *D. neb* and the two species found in subgenus *Sophophora* (Fig. 4H). Thus, the *nos* and *pgc* composition found within germ granules does not fall within the expected phylogenetic relationship. As other mRNAs and species are included from additional studies, it is possible that a more accurate phylogenetic relationship will emerge. Alternatively, the rapid evolution of 3’ UTRs has been implicated as an important genomic meta-regulator that has contributed to the adaptation, diversification, and speciation of cichlid fishes [38]. Thus, it may be that the noncoding genetic components that participate in germ granule assembly, such as 3’ UTRs, are diverging relatively quickly, producing an unexpected phylogenetic picture when comparing germ plasms due to functional impacts on germ granule compositions. Regardless, we provide evidence that the evolutionary divergence of 3’ UTRs can influence germ granule mRNA compositions. This discovery sparks curiosity about the wider influence that evolution has on biomolecular condensate structures, the fine-tuning of condensate function, and whether condensate diversity offers natural selection advantages. Further investigation is needed to explore these possibilities. The goal of this initial study was to shed light on how evolution impacts the development and composition by establishing the presence of germ granule diversity within genus *Drosophila* and identifying mechanisms that contribute to the natural diversification of germ granules. Additional classes of biomolecular condensates control various cell processes, including stress response, neurological function, cell signaling, and gene regulation [39–45]. Thus, additional studies should explore whether the mechanisms described here can be applied to other systems to gain additional knowledge regarding the evolution and development of biomolecule condensates.

## Material and Methods

### smFISH and Microscopy

Custom species specific smFISH probes were designed using Biosearch Technologies Stellaris Probe Designer tool and purchased from Biosearch Technologies where all *pgc* probe sets were labeled with Quasar 570 dye and all *nos* probe sets were labeled with ATTO 647N dye. Probe designs for *pgc* were based on the full transcript length *D. mel* (FlyBase, FBgn0016053), UniProt entries A0A6I8V605 for *D. pse*, and B4LMX3 for *D. vir*. Probe designs for *nos* were based on the coding sequence for all species, FBgn0002962 for *D. mel* (FlyBase), UniProt entry A0A6I8V5B3 for *D. pse*, and Q24710 for *D. vir*. For *D. neb*, the *nos* and *pgc* sequences were identified using the *D. neb* annotated long-read sequenced genome [46]. For *D. mel, cycB*, and *gcl* probes targeting their coding sequence were generated based on FlyBase sequences FBgn0000405, labeled with CAL Fluor Red 590 dye, and sequence FBgn0005695, labeled with CAL Fluor Red 610 Dye, respectively. smFISH experiments were performed as previously described [17, 19, 47]. In summary, females were fed yeast for 24 hours and ovaries were dissected in cold PBS. For embryo collections, flies were placed in fly cages and allowed to lay eggs for 1 hour on apple juice agar plates with yeast paste. Eggs were collected and 0-1 hour old embryos were dechorionated and devitellinized as previously described [47]. Tissue fixation was carried out for 30 minutes in 4% paraformaldehyde in PBS. All samples were mounted in Prolong Glass (Life technologies) and allowed to cure for 3 days prior to image acquisition [17, 47]. Confocal microscopy was performed as described in detail [19] using a Leica STELLARIS 5 white light laser system in photon counting mode while super-resolution images were acquired using Leica LIGHTNING deconvolution.

### Single mRNA Identification and Homotypic Cluster Quantification

Identification and quantification of single transcript RNPs and homotypic clusters were performed using the custom MATLAB (Mathworks) program used in previous germ granule studies [17, 20, 48]. In brief, we use a user defined polygon to mark the entire germ plasm within a z stack (~5μm,15 confocal slices). Next, quantification of the number of transcripts within a homotypic cluster was calculated by first setting an intensity threshold based on the average intensity of single transcript RNPs found outside of the posterior germ plasm and normalizing the intensity of germ plasm RNPs to the average intensity of single transcript RNPs [17]. To determine whether homotypic clusters of *nos* and *pgc* reside within the same germ granule (referred to as co-localized) two previously established criteria were used 1) the two homotypic clusters must be within a z distance of two slices for confocal images; and 2) they must be within a distance cutoff of 200 nm in x-y [17, 19]. Following the identification and locations of *nos* and *pgc* clusters, the data were used to create the Granule Census as previously described in detail [19, 20]. All confocal images shown in the figures were filtered by a balanced circular difference-of-Gaussian with a center radius size of 1.2 pixels and surround size of 2.2 pixels [17]. To determine that single transcript RNPs diffuse through the bulk cytoplasm in non-*D. mel* species, each species’ smFISH experiments were first imaged under identical experimental and imaging conditions as *D. mel* samples, including using the same number of probes and dyes within a *nos* or *pgc* probe set. Next, single transcript thresholds from *D. mel* smFISH experiments were applied to *nos* or *pgc* RNPs found in the bulk cytoplasm’s of non-*D. mel* species, producing comparable distributions between species where ~85% of RNPs within the bulk cytoplasm were identified as singles (Supp Fig. 1A,B). Some diffusing RNPs were calculated to have more than 1 transcript due to the random overlapping within the bulk cytoplasm that has been previously documented [17].

### Quantification of Transcript levels

For all species, females were fed yeast for 24 hours and stage 13/14 oocytes were dissected in cold PBS and isolated from the ovaries. Oocytes were homogenized and RNA was extracted using RNeasy kit (Qiagen, cat #: 74104) using the provided protocol. Following RNA extraction, the QuantiTect Reverse Transcription kit (Qiagen, cat #: 205311) was used to synthesize cDNA. For all species, qPCR experiments were performed using TaqMan Gene Expression Assays (ThermoFisher). In *D. mel, nos* (Assay ID: Dm02134535_g1), *rpl7* (Assay ID: Dm01817653_g1), and *osk* (Assay ID: Dm02134538_g1), and a custom assay that was previously designed for *pgc* were used [20]. All assays were carried out using TaqMan assay master mix (cat #: 4369514) using the included standard TaqMan protocol which was performed with a BIO-RAD CFX96 Real-Time System. Three technical replicates for three biological replicates were performed for all qPCR experiments. Each biological replicate included >15 stage13/14 oocytes that were collected from multiple females. For non-*D.mel* species, the custom TaqMan probe design tool (ThermoFisher) was used to design custom assays for *nos* and *pgc* using the same sequences as described for smFISH probe designs. For *rpl7* sequences, UniProt entry numbers B4M9C2 and Q29NI0 were used for *D. vir* and *D. pse*. For *osk* sequences, UniProt entry numbers B4LXK5 and A0A6I8URE4 were used for *D. vir* and *D. pse*. For D. *neb*, *rpl7* and *osk* sequences were determined using the *D. neb* annotated long-read sequenced genome [46]. For all expression assays, the presented fold gene expression levels are relative to *D. mel* and were calculated using the 2(-Delta Delta C(T)) method [49] using *rpl7* as an internal control [20].

### Graphing and Statistical Analyses

All Granule Censuses and heatmaps were created using MATLAB (Mathworks) as previously described [19]. GCTAnalysis scores were calculated by first subtracting two Granule Censuses to create a new matrix that represents the difference between a census pair. Next, we calculated the magnitude of the new matrix using the norm function in MATLAB (Mathworks), resulting in a value that denotes the magnitude of change between a set of Granule Censuses. Additional graphs were generated using R statistical programming, RStudio and the ggplot2 package [50–52]. The Venn diagram was created using the eulerr R package [53], undirected network graphs were created using the igraph R package [54], the density plots and their percent overlap values were calculated using the overlapping R package [55, 56]. One-way ANOVA with Dunnett’s post hoc tests were performed using the DescTools R package [57] to determine statistical significance when comparing average values from each species to *D. mel*.

### Fly Strains and Cloning

The *y^1^, w^67c23^* strain, Bloomington *Drosophila* Stock Center [BDSC] 6599, was used as *D. mel*. To label the germ granule protein ensemble in Fig. 1 the *osk-gfp* transgene (fTRG_1394), a gift from H. Jambor, was used [58]. *D. virilis, D. pseudoobscura*, and *D. nebulosa* (gifts from N. Yakoby [37, 59]), were used in this study. The *D. mel ^neb nos^* ^3’ UTR^ and *D. mel ^pse nos^* ^3’ UTR^ transgenes were created by first isolating species specific *nos* 3’ UTR from genomic DNA using PCR with primers with EcoRI and XhoI cut sites engineered at the ends of the forward and reverse primers and amplified with Phusion DNA Polymerase (NEB). Primers were designed based on the *D. pse nos* sequence used to designed the smFISH probe and the *nos* sequence for *D. neb* was identified from a long-read sequenced genome [46]. Next, we removed the *nos* 3’ UTR from a 4.3-kb *D. mel* genomic *nos* rescue fragment (a gift from the Gavis Lab [60]), using EcoRI and XhoI (NEB) and ligated species specific 3’ UTR with T4 DNA ligase (NEB). Each of the full length chimeric *nos* rescue fragments were cloned into pattB vectors using NotI (NEB) cut sites and inserted into the attP40 landing site by phiC31-mediated recombination [61]. Injection services were provided by Rainbow Transgenics. The chimeric *nos* transgenes were introduced into *nos^BNX^* homozygous females to create the *D. mel ^neb nos^* ^3’ UTR^ and *D. mel ^pse nos^* ^3’ UTR^ fly lines that only express chimeric *nos* mRNA, as similarly done previously [20]. All fly stocks, including non-*D. mel* species, were maintained at 23°C on standard cornmeal food (Bloomington formulation, Genesee Scientific Cat #: 66-112). To validate chimeric sequences, constructs were sequenced by Eton Biosciences with a *nos* primer that binds near the 3’ end of *nos* coding region, GCGATCAAGGCGGAATCGTTCC.

### Computational Modeling and Sequence Analysis

Computational modeling was carried out as previously described [20] by adjusting the model to recapitulate each species’ biological data. Clustering factor values were determined as previously described [20]. In summary, qPCR data was first applied to the model’s parameters to recreate known biological conditions. Next, homotypic cluster formation was modeled with different clustering values while maintaining the known biological conditions. Next, standard curves were generated based on plotting clustering factor values against the average cluster sizes that they produced. Biological averages were then fitted to the standard curve to determine the clustering factor value for the biological data. Estimates of evolutionary divergence between *nos* 3’ UTR sequences was determined using the Maximum Composite Likelihood model [62], based on the ClustalW alignment created by the 3’ UTR sequences from *D. mel, D. pse*, and *D. neb*. All ambiguous positions were removed for each sequence pair (pairwise deletion option) and there were a total of 1006 positions in the final dataset. Analyses were conducted in MEGA X [63].

## Acknowledgements

We thank the Center for Biological Imaging at Kean University for assisting with image acquisition and all the members of the Niepielko Lab for their helpful comments and fruitful discussions. We thank the Yakoby Lab for providing us with different *Drosophila* species and the Gavis Lab for stocks and reagents. We thank Kean University for supporting F.N.B. through the Graduate Research Assistantship program, undergraduates B.A., A.J.K., M.M.M, G.C.N, and A.L.S through the Research First Initiative, and M.G.N. through the Release Time for Research program. M.G.N. was supported by the Eunice Kennedy Shriver National Institute of Child Health & Human Development of the National Institutes of Health under Award Number R15HD102960 and by the National Science Foundation CAREER Award 2237390, co-funded by the Genetic Mechanisms Program in the Molecular and Cellular Biosciences Division and the Developmental Systems Program in the Integrative Organismal Systems Division.

## Author Contributions

Conceptualization: D.A.D, F.N.B., and M.G.N.; Methodology and Investigation: D.A.D, F.N.B., B.A., A.J.K., M.M.M., G.C.N, B.U.P., G.A.H., A.L.S, C.J.S., and M.G.N.; Software: B.M.O. and M.G.N.; Writing, review and editing: D.A.D, F.N.B., and M.G.N.; Supervision: M.G.N.; Funding acquisition: M.G.N.

## Declaration of Interests

The authors declare no competing interests.

## Inclusion and Diversity

We support inclusive, diverse, and equitable conduct of research and strive to advance these values in all our activities as members of New Jersey’s Research Alliance for Inclusive STEM Education (NJ-RAISE).

## Notes

### Competing Interest Statement

The authors have declared no competing interest.

