## Supplement Figures 1-4 for "Evolutionary changes in germ granule mRNA content are driven by multiple mechanisms in *Drosophila*"

**Supplement Figure 1**

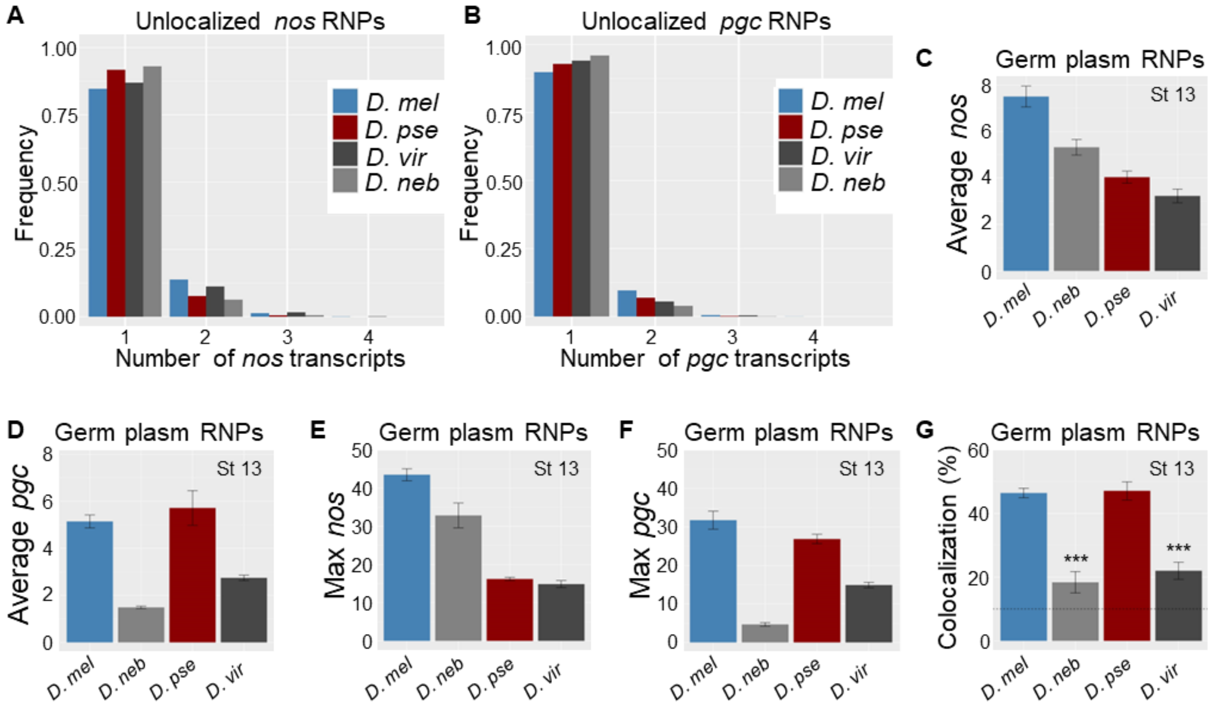


**Supplement Figure 1: Number of *nos* and *pgc* transcripts in RNPs from the bulk oocyte cytoplasm and germ plasm. A)** Distribution of the number of *nos* transcripts found in unlocalized RNPs in the bulk oocyte cytoplasm in all species analyzed n > 10,000 RNPs. **B)** Distribution of the number of *pgc* transcripts found in unlocalized RNPs in the bulk oocyte cytoplasm in each species tested n > 10,000 RNPs. **C)** Average number of *nos* transcripts found in *nos* containing germ plasm RNPs found in each species in stage 13 oocytes. **D)** Average number of *pgc* transcripts found in *pgc* containing germ plasm RNPs found in each species in stage 13 oocytes. **E)** Largest number of transcripts (Max) observed in *nos* containing germ plasm RNPs for each species in stage 13 oocytes. **F)** Max number of *pgc* transcripts found in *pgc* containing germ plasm RNPs for each species in stage 13 oocytes. **G)** Colocalization rate between *nos* and *pgc* containing RNPs in the germ plasm for each species in stage 13 oocytes. For C-G n > 5,000 germ plasm RNPs for each mRNA type. All error bars represent S.E.M. based on data collected from n at least 3 stage 13 oocytes. The dotted horizontal line represents the expected rate for random co-localization.

### Supplement Figure 2

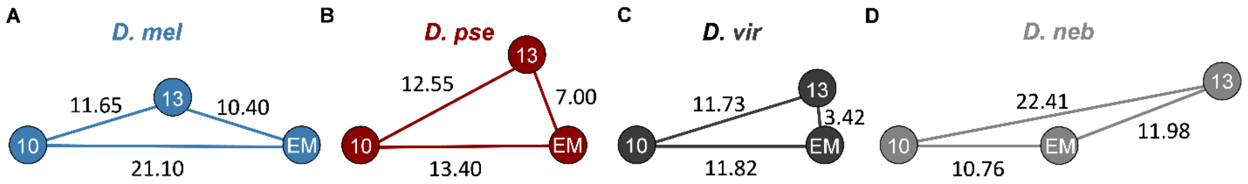


**Supplement Figure 2: Representation of germ granule assembly dynamics using GCTAnalysis scores. A-D)** Visual representation of germ granule formation dynamics where each edge represents a GCTAnalysis score that was calculated between each developmental stage, represented as nodes.

### Supplement Figure 3

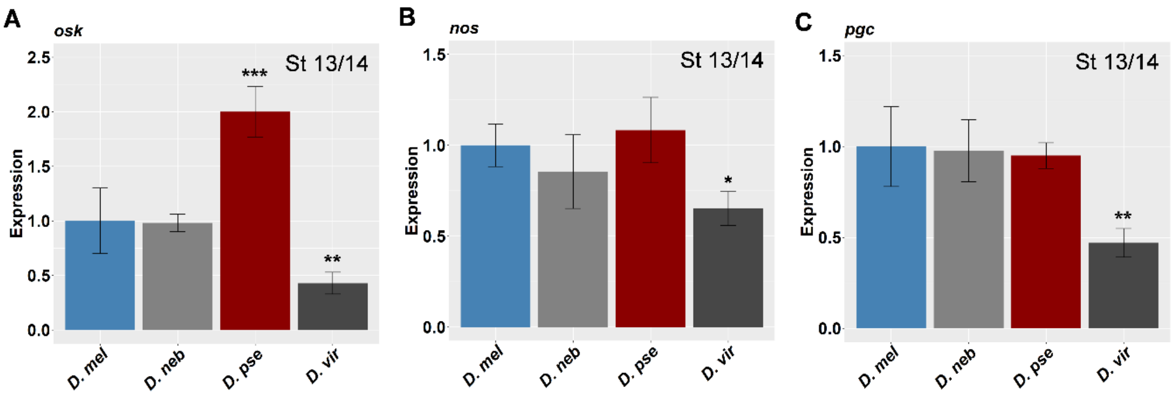


**Supplement Figure 3: Relative mRNA expression levels of *osk, nos*, and *pgc* in different *Drosophila* species. A-C)** Relative mRNA expression levels for **A)** *osk*, **B)** *nos*, and **C)** *pgc*. All expression levels are relative to *D. mel* and were measured from stage 13/14 oocytes. Error bars represent S.E.M. from n = 3 biological replicates.

**Supplementa Figure 4**

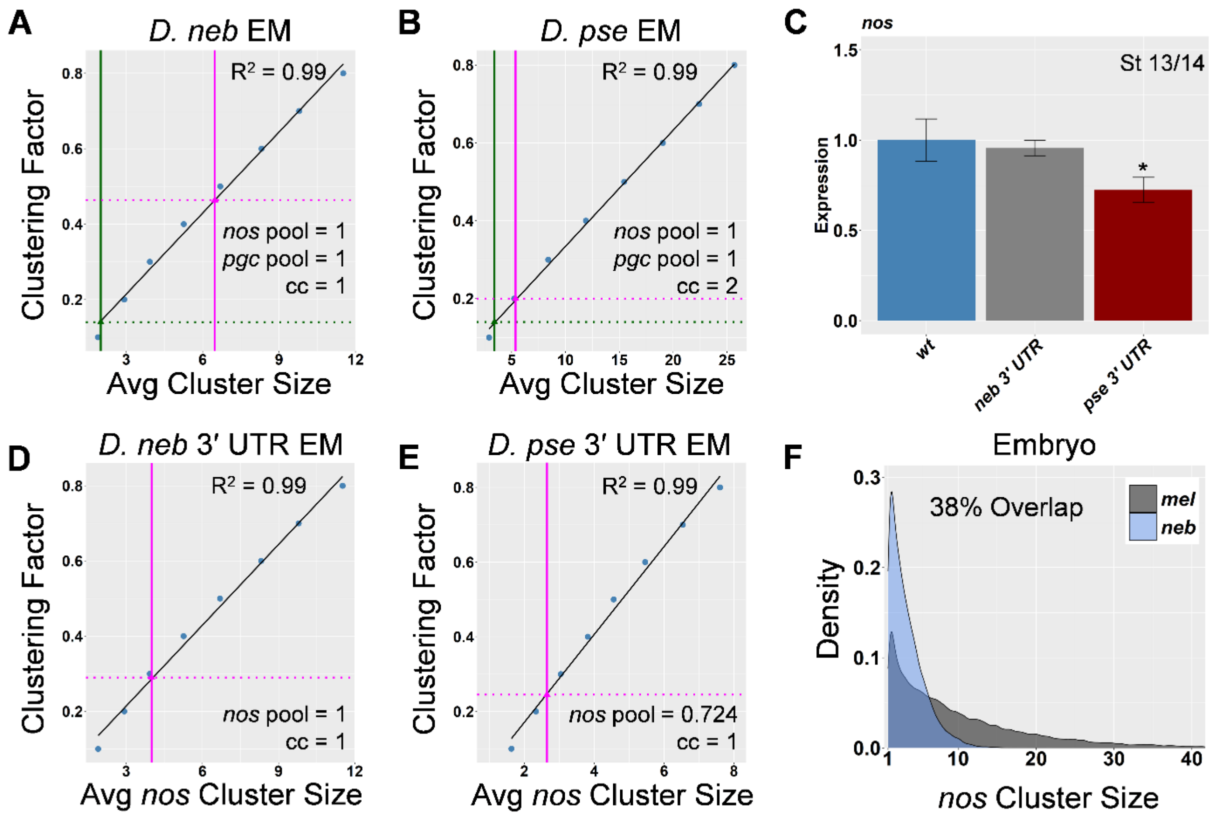


**Supplement Figure 4: Calculating the clustering factor parameter, relative *nos* expression levels from recombinant flies, and *nos* cluster size density plots. A-B)** Standard curves produced by the computational model using the biological determined parameters for *nos* expression (*nos* pool), *pgc* expression (*pgc* pool), and *osk* expression which represents carrying capacity (cc). The standard curves (solid black line) are produced using the average cluster size (x-axis) that is produced for a given clustering factor (y-axis). Fitting the biologically determined average cluster size for *nos* (vertical magenta line) and average cluster size for *pgc* (vertical green line) reveals the clustering factor for *nos* (broken horizontal magenta line) and for *pgc* (broken horizontal green line) for **A)** *D. neb* and **B)** *D. pse* embryos. **C)** Relative *nos* expression levels from recombinant *D. mel* flies that contain the *D. neb* or the *D. pse nos* 3′ UTR. Expression levels are relative to wild type *D. mel nos* levels and were measured from stage 13/14 oocytes. Error bars represent S.E.M, n = 3 biological replicates. **D-E)** Standard curves produced by the computational model using the biological determined parameters for *nos* expression (*nos* pool) and *osk* expression which represents carrying capacity (cc). The standard curves (solid black line) are produced using the average *nos* cluster size (x-axis) that is produced for a given clustering factor (y-axis). Fitting the biologically determined average cluster size for *nos* (vertical magenta line) reveals the clustering factor for *nos* (broken horizontal magenta line) for **D)** *D. neb nos* 3′ UTR in *D. mel* and **E)** *D. pse nos* 3′ UTR in *D. mel*. **F)** Density plots for *nos* cluster sizes in the embryo for wild-type *D. mel* (gray) and *D. mel* that contains the *nos* 3′ UTR from *D. neb* (blue), plots were generated using >17,500 *nos* clusters for each genotype from at least 4 germ plasms.
